# Comparative genomic analysis of the emerging pathogen *Streptococcus pseudopneumoniae*: novel insights into virulence determinants and identification of a novel species-specific molecular marker

**DOI:** 10.1101/468462

**Authors:** Geneviève Garriss, Priyanka Nannapaneni, Alexandra S. Simões, Sarah Browall, Raquel Sá-Leão, Herman Goossens, Herminia de Lencastre, Birgitta Henriques-Normark

**Author notes:** These authors contributed equally (Geneviève Garriss, Priyanka Nannapaneni). Corresponding author: Birgitta Henriques-Normark, Professor, MD, Karolinska University hospital and Karolinska Institutet, MTC, Nobels väg 16, SE-171 77 Stockholm, Sweden.

## Abstract

*Streptococcus pseudopneumoniae* is a close relative of the major human pathogen *S. pneumoniae*. While initially considered as a commensal species, it has been increasingly associated with lower-respiratory tract infections and high prevalence of antimicrobial resistance (AMR). *S. pseudopneumoniae* is difficult to identify using traditional typing methods due to similarities with *S. pneumoniae* and other members of the mitis group (SMG). Using phylogenetic and comparative genomic analyses of SMG genomes, we identified a new molecular marker specific for *S. pseudopneumoniae* and absent from any other bacterial genome sequenced to date. We found that a large number of known virulence and colonization genes are present in the core *S. pseudopneumoniae* genome and we reveal the impressive number of known and new surface-exposed proteins encoded by this species. Phylogenetic analyses of *S. pseudopneumoniae* show that specific clades are associated with allelic variants of core proteins. Resistance to tetracycline and macrolides, the two most common resistances, were encoded by Tn*916*-like integrating conjugative elements and Mega-2. Overall, we found a tight association of genotypic determinants of AMR as well as phenotypic AMR with a specific lineage of *S. pseudopneumoniae*. Taken together, our results sheds light on the distribution in *S. pseudopneumoniae* of genes known to be important during invasive disease and colonization and provide insight into features that could contribute to virulence, colonization and adaptation.

**Importance:** *S. pseudopneumoniae* is an overlooked pathogen emerging as the causative agent of lower-respiratory tract infections and associated with chronic obstructive pulmonary disease (COPD) and exacerbation of COPD. However, much remains unknown on its clinical importance and epidemiology, mainly due to the lack of specific means to distinguish it from *S. pneumoniae*. Here, we provide a new molecular marker entirely specific for *S. pseudopneumoniae*. Furthermore, our research provides a deep analysis of the presence of virulence and colonization genes, as well as AMR determinants in this species. Our results provide crucial information and pave the way for further studies aiming at understanding the pathogenesis and epidemiology of *S. pseudopneumoniae*.

## Introduction

*Streptococcus pseudopneumoniae* is a close relative of the human pathogen *Streptococcus pneumoniae*. It was first described in 2004 (1), and belongs to the mitis group which includes 13 other species of which some are the most common colonizers of the oral cavity, such as *S. mitis*, *S. sanguinis*, *S. oralis* and *S. gordonii* (2). An increasing number of reports indicate that *S. pseudopneumoniae* is a potential pathogen, usually associated with underlying conditions (3-5), and that it can be isolated from both invasive and non-invasive sites (6-9). It has been shown to be virulent in a mouse peritonitis/sepsis model (10), and to be the probable causative agent of fatal septicemia cases (5). Rates of antimicrobial resistance (AMR) have been reported to be high in several studies, in particular to penicillin, macrolides, co-trimoxazole and tetracycline (6-8). However, despite its emergent role as a pathogen, relatively little is known on its epidemiology, pathogenic potential and genetic features.

Recent studies revealed that more than 50% of the publicly available genome sequences of *S. pseudopneumoniae* strains in fact belong to other species of the mitis group (11, 12), highlighting the challenges faced when identifying strains of this species. *S. pseudopneumoniae* was originally described as optochin-resistant if grown in presence of 5% CO_2_, but susceptible in ambient atmosphere, bile insoluble and non-encapsulated (1). Exceptions to these phenotypes were later reported (4, 5, 7, 13). Several molecular markers previously thought to be specific for *S. pneumoniae*, such as 16S rRNA, *spn9802*, *lytA*, *ply* and *pspA*, have been used in PCR-based assays, but were subsequently discovered in some *S. pseudopneumoniae* isolates (7, 13, 14). In addition, the inherent problem of these markers is that they aim at identifying pneumococci and thus have limited value for the positive identification of *S. pseudopneumoniae*. To date, only one molecular marker has been described for the identification of *S. pseudopneumoniae*, however it is found in a subset of *S. pneumoniae* strains (12). Multi-locus sequence analysis (MLSA) is currently considered as the gold standard; however it faces limitations, such as the lack of amplification of certain alleles, or because certain isolates fail to fall within a specific phylogenetic clade (7). Understanding the clinical significance and epidemiology of *S. pseudopneumoniae* requires more discriminative identification methods and more complete picture of its genetic diversity.

The polysaccharide capsule is one of the major virulence factors of *S. pneumoniae*, due to its inhibitory effect on complement-mediated opsonophagocytosis, however a plethora of other factors and especially surface-exposed proteins have been shown to significantly contribute to pneumococcal disease and colonization (reviewed in (15, 16)). Despite the lack of a capsule, naturally non-encapsulated pneumococci (NESp) can cause disease, in particular conjunctivitis and otitis media (reviewed in (17)). The pneumococcal surface protein K (PspK) expressed by a subgroup of NESp has been shown to promote adherence to epithelial cells and mouse nasopharyngeal colonization to levels comparable with encapsulated pneumococci (18, 19), pointing to the advantage that surface-exposed proteins might provide to non-encapsulated strains.

Some studies have described the presence of pneumococcal virulence genes in *S. pseudopneumoniae* (3, 9, 20, 21), but a comprehensive overview of the distribution of known, and potentially new, genes that could promote virulence and colonization in this species is lacking.

The aim of this study was to use phylogenetic and comparative genomic analyses to identify a new molecular marker for the specific identification of *S. pseudopneumoniae* and to analyse the distribution of known pneumococcal virulence and colonization factors in this species. In addition, we have found a tight association of AMR with certain lineages and uncovered a large number of novel surface-exposed proteins.

## RESULTS

### Identification of *S. pseudopneumoniae* genomes

A first phylogenetic analysis, including 147 genomes from various streptococci of the mitis group (SMG) species, was performed to classify 24 isolates collected from lower-respiratory tract infections (LRTI) within the EU project GRACE (22) which we suspected to be *S. pseudopneumoniae* (n=16) or *S. mitis* (n=3), or for which no definitive classification was possible to obtain using traditional typing methods and MLSA (n=5) (Fig. 1). 21/24 LRTI isolates clustered within the *S. pseudopneumoniae* clade, including the strains for which a precise MLSA identification had not been possible to obtain. The 3 strains initially identified as *S. mitis* clustered within the *S. mitis* clade and are not discussed further. In line with earlier observations (11, 12), 8 non-typable *S. pneumoniae* genomes fell within the *S. pseudopneumoniae* clade, along with only 15/38 publicly available genomes currently classified as *S. pseudopneumoniae* (Fig. 1). Based on our phylogenetic analysis, a total of 44 sequenced genomes were considered as *S. pseudopneumoniae* and further analyzed (Table S1).

**Fig. 1.**
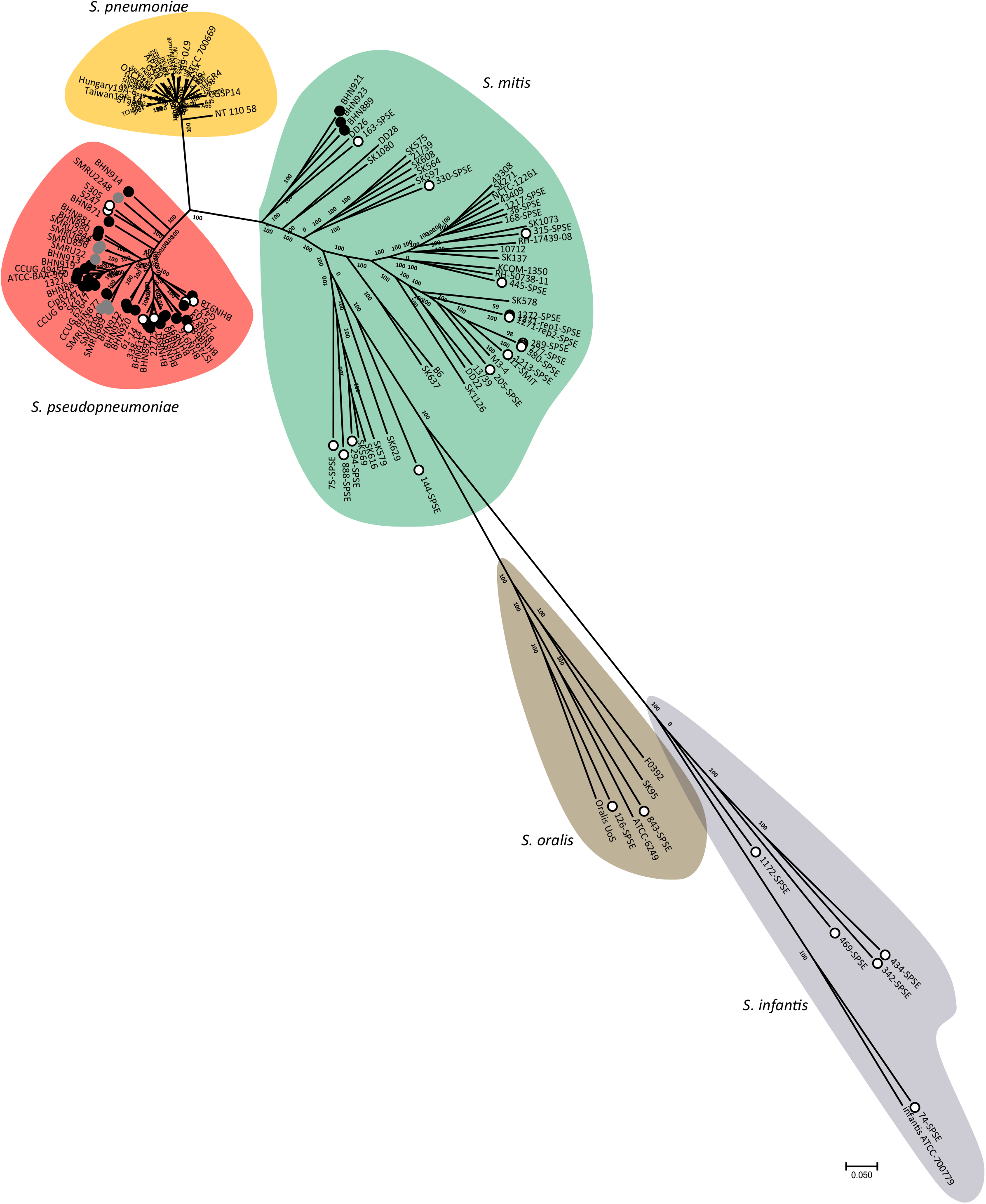
SMG unrooted consensus parsimony phylogenetic tree based on all SNPs (1230968) of 147 genomes: LRTI isolates (24) and publicly available *S. pseudopneumoniae* (38), *S. pneumoniae* (39), NT *S. pneumoniae* (8), *S. mitis* (36), *S. oralis* (1) and *S. infantis* (1). Circles indicate isolates from lower-respiratory tract isolates (black), NCBI genomes labelled as *S. pseudopneumoniae* (open) or NT *S. pneumoniae* (grey). Background shading delineates clades of different species. The tree was built in kSNP and visualized in MEGA7 (50).

### A single gene, SPPN_RS10375, can be used to identify *S. pseudopneumoniae*

In the course of the initial characterization of the LRTI isolates, we observed that 13/21 displayed the typical optochin susceptibility and bile solubility phenotypes previously attributed to *S. pseudopneumoniae* (1) (Table 1). Using the whole genome sequencing (WGS) data, we sought to clarify the discrepancy between the RFLP and PCR results used for detecting the pneumococcal variant of *lytA*. This revealed that some *S. pseudopneumoniae* phage-encoded *lytA* genes could be similar enough to be detected by PCR as the pneumococcal *lytA*, but that they lacked the BsaAI restriction site used for RFLP analysis (Fig. S1) (14). In addition, the pneumococcal variant of *ply* was detected by RFLP in three instances, but we found that while these genes cluster in separate clade, they harbor the restriction site used for RFLP identification of pneumococcal *ply* (Fig. 2) (7, 9).

**Fig. 2.**
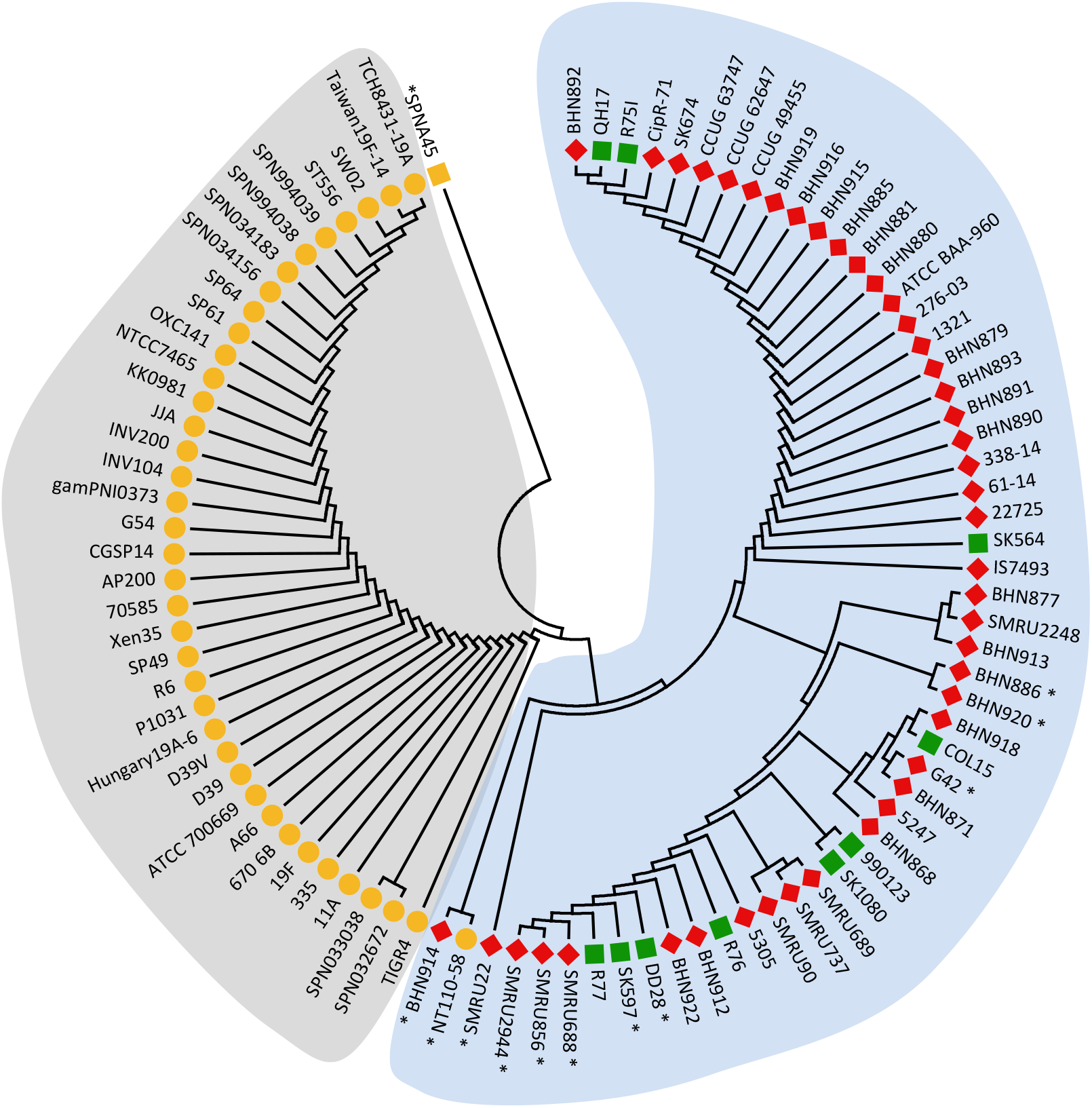
Phylogenetic tree of 93 Ply alleles from SMG species. MEGA7 (50) was used to infer the evolutionary history using the Maximum Likelihood method based on the JTT matrix-based model (55). The tree with the highest log likelihood (−1453.23) is shown. There were a total of 245 positions in the final dataset. Leafs are colored based on the species: yellow, *S. pneumoniae*; red, *S. pseudopneumoniae*; green, *S. mitis*, and Ply clades are indicated by the background shading: grey, pneumococcal Ply; blue, atypical (Mly/Pply). Asterisks indicate Ply variants outside of the *S. pneumoniae* Ply clade that would be classified as pneumococcal Ply based on the presence of the BsaAI restriction site used for RFLP analysis.

**Table 1.** Phenotypic and genotypic characterization of LRTI isolates belonging to the *S. pseudopneumoniae* phylogenetic clade.

|  | Number of strains<br>(n=21) | % |
| --- | --- | --- |
| **Phenotypic markers** |  |  |
| Optochin susceptibility |  |  |
| 5% CO<sub>2</sub> | 2 | 9,5 |
| Ambient atmosphere | 14<sup>a</sup> | 66,7 |
| Bile solubility | 1<sup>b</sup> | 4,8 |
| **Genotypic markers** |  |  |
| PCR markers |  |  |
| Pneumococcal *lytA* | 2 | 9,5 |
| *cpsA* | 0 | 0,00 |
| *spn9802* | 20 | 95,2 |
| Pneumococcal-specific 16S rRNA | 19 | 90,5 |
| RFLP signatures |  |  |
| Pneumococcal/atypical *lytA* | 0/21 | 0/100 |
| *ply/mly* | 3/18 | 14,3/85,7 |

<sup>a</sup>7 strains did not grow in ambient atmosphere. The 14 strains susceptible in ambient atmosphere were resistant in CO<sub>2</sub>.
<sup>b</sup>2 strains showed partial solubility.

We sought to identify a single genetic locus uniquely present in all strains of the *S. pseudopneumoniae* clade. We determined the pan genome of *S. pseudopneumoniae* and *S. pneumoniae* and identified 30 clusters of orthologous genes (COGs) present in the 44 *S. pseudopneumoniae* genomes, but absent from the 39 *S. pneumoniae* completed genomes (Table S2). BLAST analysis revealed that SPPN_RS10375 and SPPN_RS06420 were not found in any genome belonging to other species but *S. pseudopneumoniae*. While SPPN_RS06420 had a G+C content challenging for the design of PCR primers (average of 27.1%) further analysis of SPPN_RS10375 and its surrounding intergenic regions in the 44 genomes indicated that this 627-bp locus could be a good candidate for a molecular marker. 8 clinical isolates, not subjected to whole-genome sequencing, and collected during the same LRTI study (22), that were either impossible to identify (n=4) or suspected to be *S. pseudopneumoniae* (n=4), were found to be positive by PCR for SPPN_RS10375, indicating they are all *S. pseudopneumoniae*. These strains were also positive for the recently published *S. pseudopneumoniae* marker SPS0002 (12) (Fig. S2).

### Pan and core genome analyses of *S. pseudopneumoniae*

The closest relative of *S. pseudopneumoniae* is *S. pneumoniae*, however, no study has yet investigated in depth the genetic similarities and differences that characterize them. We defined the pan-genome of these two species using the 44 *S. pseudopneumoniae* genomes and 39 completed and fully-annotated *S. pneumoniae* NCBI genomes (Table S3). 1236/4548 COGs (27%) were unique to *S. pseudopneumoniae*, while 1126 (25%) were unique to the pneumococcus. The remaining 2186 COGs (48%) were shared by both species. To evaluate the presence in *S. pseudopneumoniae* of infection/colonization relevant genes, we investigated the presence of 356 *S. pneumoniae* genes differentially expressed in mice models of invasive disease and during epithelial cell contact (23), and found that 94% are present in at least one *S. pseudopneumoniae* genome (Table S4). 74% of these genes were found in the core genome of the 39 completed *S. pneumoniae* genomes (100% of the genomes). While fewer (53%) of these genes were found in the core *S. pseudopneumoniae* genome, the use of draft *S. pseudopneumoniae* genomes in contrast with fully assembled *S. pneumoniae* genomes likely results in an underestimation of their presence. 20/356 genes were absent from *S. pseudopneumoniae* and amongst them was the gene encoding pneumococcal surface protein A (*pspA*), a known virulence factor. 8/20 absent genes are core *S. pneumoniae* genes, 4 of which are organized in an operon involved in stress response (SP_RS08945-SP_RS08960). The other 4 genes encode a product of unknown function (SP_RS11915), a putative methyltransferase (SP_RS07780) and two products predicted to be involved in co-factor metabolism (SP_RS10205 and SP_RS10210).

Surprisingly, results from this screen revealed that the capsular genes *cps4A* and *cps4C* (also named *wzg* and *wzd* (24)) are found in one *S. pseudopneumoniae* strain. Further analysis revealed that BHN880 harbors a capsular locus similar to pneumococcal serotype 5 and to the capsule loci of *S. mitis* strain 21/39 (Fig. 3). Gel diffusion assays typed BHN880 as pneumococcal serotype 5, which is supported by the high nucleotide identity (97.7%) between the regions encoding the sugar precursors of the BHN880 and the serotype 5 capsular loci. The 43 remaining genomes carry an NCC3-type capsule locus (19) which encompasses genes *dexB*, *aliD* and *glf* (also known as *cap* or *capN* (19, 25)).

**Fig. 3.**
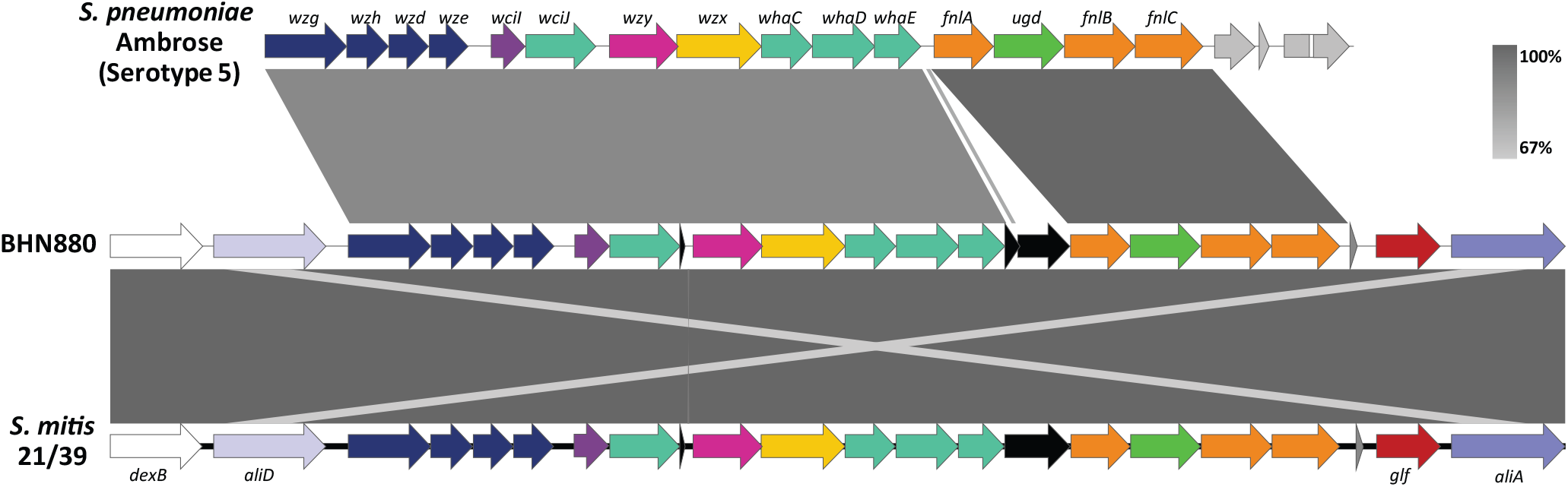
Pairwise alignment of the capsule locus of *S. pseudopneumoniae* strain BHN880 with *S. pneumoniae* Ambrose and *S. mitis* 21/39. Colors and annotations are based on Bentley *et al* (24). Grey shading indicates degree of pairwise nucleotide identity.

### Pneumococcal virulence and colonization genes are widely distributed in *S. pseudopneumoniae*

To gain greater insight into genetic features that could promote adhesion, virulence and colonization we investigated the presence of orthologues of 92 pneumococcal surface-exposed proteins, transcriptional regulators and two-component signal transducing systems (TCSs), for which the distribution among pneumococcal genomes has been studied (26, 27). Due to the fact that 43/44 genomes are draft genomes we considered proteins present in 42 of the 44 genomes to be present in all strains (core genome). 16/92 proteins had no orthologs in *S. pseudopneumoniae*, including the subunits of both pili (RrgABC and PitAB), surface-exposed proteins PsrP and PspA, and the stand-alone regulators MgrA and RlrA (Table S5). 3 of these 16 proteins, HysA, PclA and MgrA, are core *S. pneumoniae* features (26). Other core *S. pneumoniae* proteins were represented in only a very small subset of *S. pseudopneumoniae* strains, such as Eng (n=1), PiaA (n=1), GlnQ (n=3) and the HK and RR that constitute TCS06 (n=3). 29/61 surface-exposed proteins were found in the core *S. pseudopneumoniae* genome, including amongst others major virulence factors such as Ply, NanA and HtrA (Fig. 4A and Table S5). The NanA variant found in *S. pseudopneumoniae* shares similar domains and good similarity with pneumococcal NanA, however it differs strongly in its C-terminal region, where the LPxTG-anchoring domain is replaced with a choline-binding domain (CBD). Pneumococcal LPxTG-anchored proteins were found to have the lowest levels of representation in *S. pseudopneumoniae*, with 12/23 being absent from all genomes. With the exception of TCS06 and HK11 all HK-RR pairs were core *S. pseudopneumoniae* proteins. 2/3 isolates encoding TCS06 also harbor a PspC-like protein in the same locus, such as is found in pneumococcal genomes. These two PspC-like proteins carry an LPxTG-anchoring domain and share limited similarity to each other (30.8%), and to their closest pneumococcal allele, PspC11.3 (32.9%) (28). The third genome encoding TCS06 carries a truncated gene encoding a PspC-like protein.

**Fig. 4.**
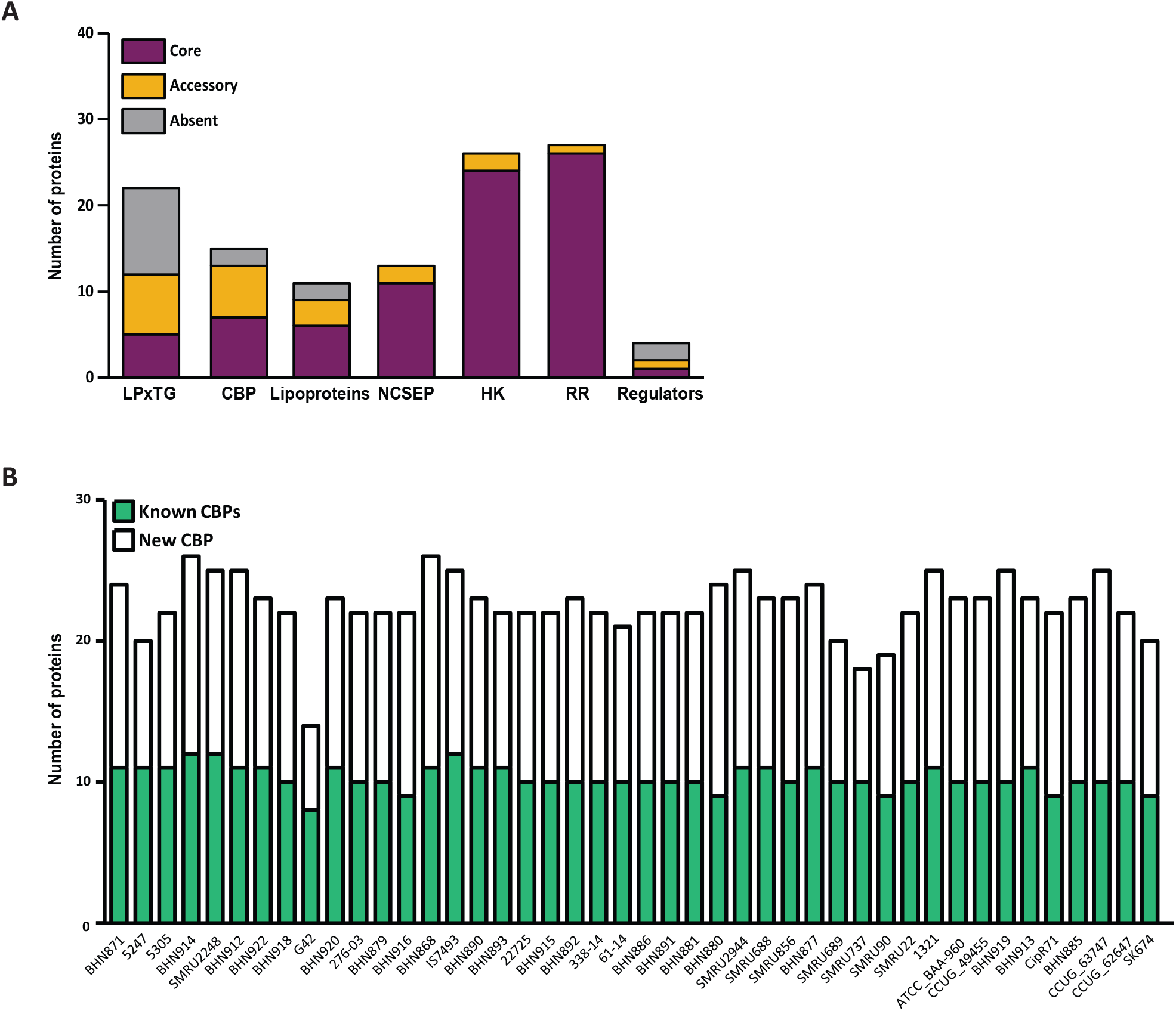
Presence of relevant pneumococcal proteins and new features in *S. pseudopneumoniae*. A) Distribution of known pneumococcal surface-exposed proteins, TCS and stand-alone regulators in *S. pseudopneumoniae*. B) Number of known and new choline-binding proteins in each *S. pseudopneumoniae* strain.

### *S. pseudopneumoniae* encodes a massive number of new surface-exposed proteins and 6 new two component systems

We then investigated if *S. pseudopneumoniae* harbored additional features that could potentially be relevant in virulence or colonization scenarios. We searched the proteome of the *S. pseudopneumoniae* species for novel choline-binding proteins (CBPs) and new TCSs. We found 19 previously undescribed proteins containing a CBD, which we named Cbp1 to Cbp19 (Table S6). 4 of these proteins belong to the core genome while the others have varying levels of presence amongst the 44 genomes. Each strain carried between 6 and 15 new CBPs, and some *S. pseudopneumoniae* genomes carried a total of 26 CBPs (Fig. 4B). The presence of signal peptides, transmembrane domains, and other know functional domains in *S. pseudopneumoniae* CBPs are summarized in Fig. 5.

**Fig. 5.**
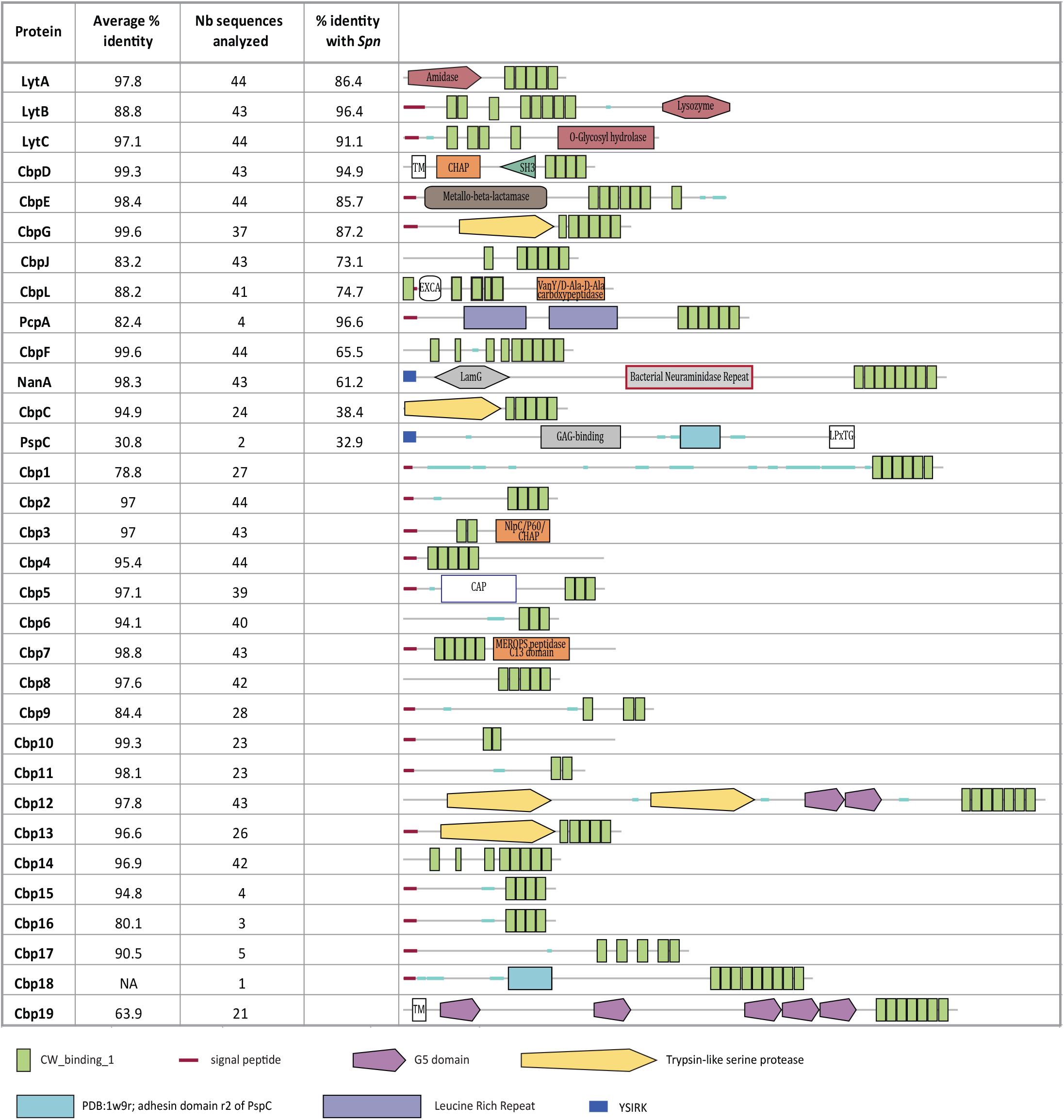
Choline-binding proteins of *S. pseudopneumoniae*. Characteristics of CBPs found in at least one *S. pseudopneumoniae* genome. Average % identity within pseudopneumoniae species and the number of proteins analyzed are indicated. % identity with *S. pneumoniae* (*Spn*) was calculated using the proteins from IS7493 and *S. pneumoniae* TIGR4, except in the following cases: NanA (R6); PspC (Allele PspC11.3-AF276622.1). Representations of domains found in each CBP are based on SMART (56) analysis of the variant found in IS7493. In absence of the protein from IS7493, analysis was based BHN914 (PspC, Cbp15, Cbp16, Cbp17, Cbp18, Cbp19); BHN879 (Cbp1) BHN886 (Cbp19).

We found six additional HK-RR pairs in the *S. pseudopneumoniae* pan-genome, 4 of which are core features (Table 2 and Table S6). We have named these TCS14 to TCS19. A more detailed analysis of their genetic loci revealed that TCS14 is found in the same loci as genes encoding a ComC/Blp family peptide and bacteriocins. These genes are distinct from the homologs of ComC and BlpC, present elsewhere in the *S. pseudopneumoniae* genome. The remaining five TCS are genetically linked to genes predicted to encode ABC transporters.

**Table 2.** Novel Two-component signalling systems of *Streptococcus pseudopneumoniae*.

| TCS | RR<sup>a</sup> | HK<sup>b</sup> | Species of closest homologue | Family of regulators | Associated genes |
| --- | --- | --- | --- | --- | --- |
| 14 | SPPN_RS00570 | SPPN_RS00565 | *S. mitis* | LytTR | Bacteriocins |
| 15 | SPPN_RS11635 | SPPN_RS01890 | *S. mitis* | YesN | Ferric iron transport |
| 16 | SPPN_RS03570 | SPPN_RS03565 | *S.pseudoporcinus*,<br>*S. canis* | OmpR | Potassium transport<sup>c</sup> |
| 17 | SPPN_RS07705 | SPPN_RS07700 | *S. parasanguinis* | LytTR/YesN | Thiamine biosynthesis<sup>d</sup> |
| 18 | BHN881_01880 | BHN881_01881 | *S. mitis* | YesN / AraC | Sugar transport |
| 19 | BHN877_00996 | BHN877_00995 | *S. suis* | CitB | Bacitracin export |

<sup>a</sup> Locus tag of the response regulator (RR). Locus in IS7493 is used when present.
<sup>b</sup> Locus tag of the sensor histidine kinase (HK). Locus in IS7493 is used when present.
<sup>c</sup> Similar to the kdpD/kdpE from *E.coli* (60).
<sup>d</sup> Similar to TCS02 of *S. thermophilus* (61).

### The most common resistances are carried by potentially mobile genetic elements

Resistance to erythromycin and tetracycline were previously reported as very common in *S. pseudopneumoniae* (4, 6-8), and they are also the two most common resistances found in our collection (Table S7). We investigated the genetic determinants encoding these resistances and found that more than half of the strains (n=24) harbored genes encoding resistance to tetracycline (*tet*(M)), 14- and 15-membered macrolides (*mef*(E)*/msr*(D)) and/or macrolides, lincosamides and streptogramin B (MLS_B_ antibiotics) (*erm*(B)). *mef*(E)*/msr*(D) genes were found to be part of a Mega-2 element (macrolide efflux genetic assembly), integrated within the coding sequence of a DNA-3-methyladenine glycosylase homolog to SP_RS00900 of *S. pneumoniae* TIGR4 (Fig. S3A). Integration of Mega-2 in this site has been previously reported in *S. pneumoniae* (29, 30). *tet*(M) and *erm*(B) genes were found within the Tn*916*-like integrating conjugative elements (ICEs) Tn*5251* (31) and Tn*3872* (32) (Fig. S3A and Table S8). Tn*5251* and Tn*3872* ICEs were highly similar between the various strains (Fig. S3B and S3C) and were found integrated in 7 different integration sites in the chromosome (Table S8). 4 of the integration sites were unique, while the other 3 were shared by two or more strains. One strain, SMRU2248, carried the *tet*(O) gene, which also encodes tetracycline resistance, in what appeared to be the remnant of a Tn*5252*-like ICE. Two strains carried an aminoglycoside-3'-phosphotransferase *aph*(3’)-Ia gene.

### Bacteriophages are tightly associated with *S. pseudopneumoniae*

27/44 *S. pseudopneumoniae* strains carried at least one putatively full-length prophage. 21 of these prophages shared a highly related integrase (≥90.5% identity nucleotide) which we termed Int*Sppn1*, and in 19 cases these prophages were found integrated between SPPN_RS05275 (encoding a putative CYTH domain protein) and SPPN_RS05395 (encoding a putative GTP pyrophosphokinase) (Table S9). The remaining 2/21 phages were found alone in a contig without chromosomal flanking sequences. Although a full-length prophage could not be confirmed in the remaining 23 strains they harbored the same integrase, which was, except in two cases (G42 and ATCC BAA_960), associated with some phage genes. 6 strains carried an additional putatively full-length phage encoding an integrase closely related to that of pneumococcal group 2a prophages (33). These prophages were found between SPPN_RS07570 and SPPN_RS07555, which are the homologs of the genes flanking the phage group 2a integration site in pneumococci (34). 23 other strains harbored this integrase, however, the presence of more than one phage per strain severely impaired our ability to confirm the completeness of the phages they were associated with, as phage sequences were split between various contigs.

### Phylogenetic clades of *S. pseudopneumoniae* are characterized by different patterns of accessory virulence genes and antibiotic resistance genes

A SNP-based phylogenetic tree using the 793 *S. pseudopneumoniae* core COGs revealed that the species is divided into three clades (Fig. 6A). Clades II and III encompass most of the isolates while clade I is composed of 5 isolates which fall closer to the *S. pneumoniae* strains (Fig. 6A and Fig. S4). All three clades were composed of strains isolated from the nasopharynx and from sputum or lower-respiratory tract samples. The three blood isolates belonged to clade II. We investigated the distribution of accessory proteins and allelic variants of core proteins in each clade, as well as the presence of genetic determinants of AMR and phenotypic resistances to penicillin and co-trimoxazole (SXT), as they had high prevalences in other reports (6-8) and were available for many of the NCBI genomes.

**Fig. 6.**
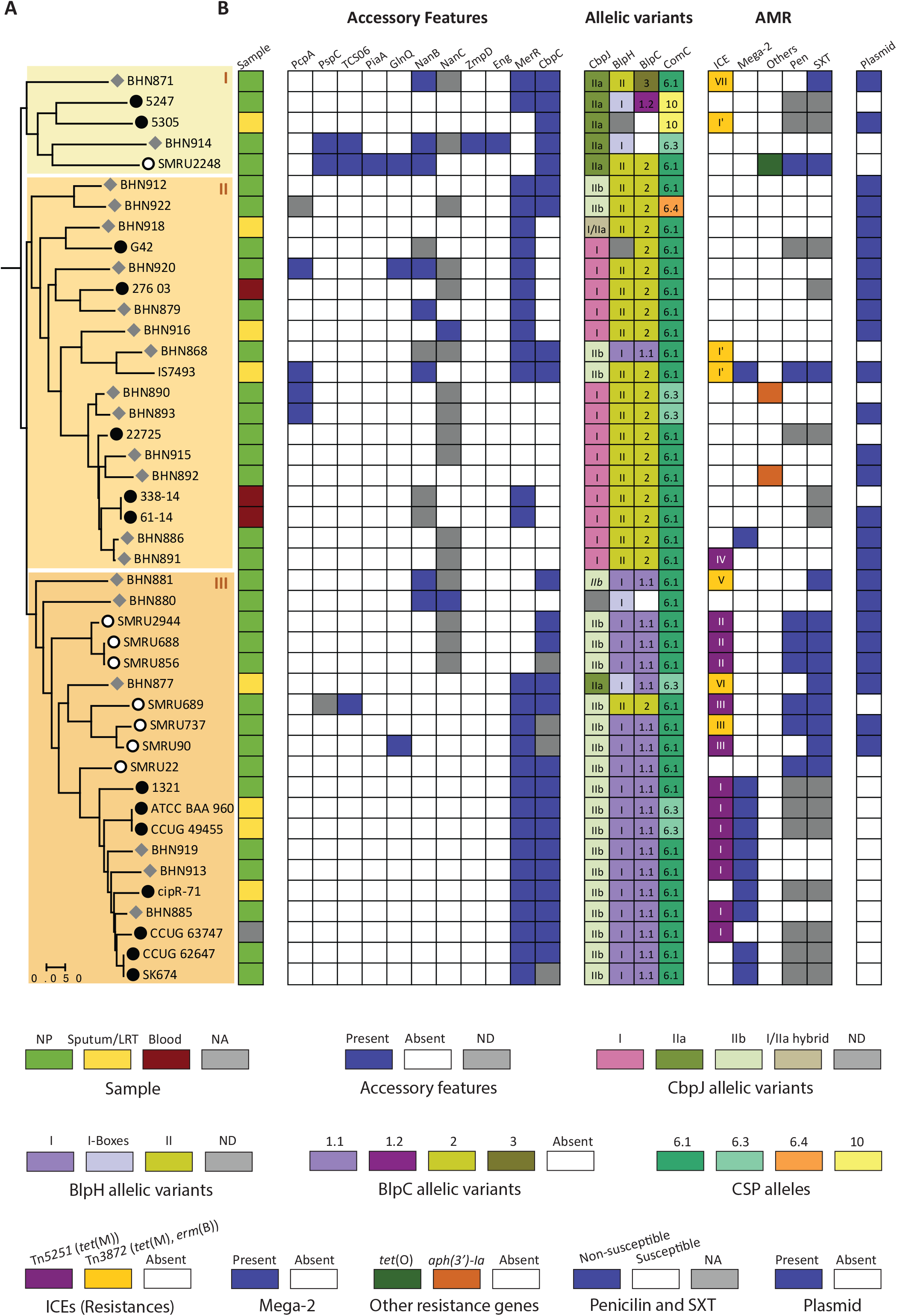
Phylogenetic distribution of accessory features and allelic variants. A) Core-genome species tree based on SNPs in 793 single copy core genes of 44 *S. pseudopneumoniae* genomes. Circles indicate isolates from lower-respiratory tract isolates (black), NCBI genomes labelled as *S. pseudopneumoniae* (open) or NT *S. pneumoniae* (grey). The tree was built in PanX and visualized in MEGA7. Clades are delineated by the background shading. B) Distribution of accessory features and allelic variants of surface exposed proteins, regulatory genes and peptide pheromones, genotypic and phenotypic antibiotic resistances, and plasmids. Description of the colors for each column is indicated in the key. Supporting information on allelic variants can be found in Fig. S5. Roman numerals in column "ICE" refers to integration sites (Table S8). ICE, Mega-2 and "other resistances" refer to genotypic resistances; penicillin (Pen) and co-trimoxazole (SXT) refer to phenotypic resistances (Table S7) and references (5, 20, 57-59). ND, not determined due to the presence of pseudogenes/contig breaks; NA, data not available.

PcpA was found exclusively in clade II, while PiaA, ZmpD and Eng were found in clade I (Fig. 6B). GlnQ, NanB and NanC were not associated with any particular clade. While MerR was well represented in all clades, CbpC was found in most strains of clade I and all strains of clade III. The presence of CbpC correlated with specific alleles of CbpJ (Fig. 6B and S5A). Strains which carried variant I of CbpJ were exclusively found in clade II and where in all cases devoid of CbpC. BlpH proteins (HK13) belonged to one of two variants which were tightly associated with clade II and clade III (Fig. 6B and S5B). Four variants of BlpH which did not specifically cluster with a specific clade were found to be similar to BlpH-I in boxes 1 and 2, which are important for interaction with BlpC (35). As expected, BlpH variants were almost strictly associated with specific variants of BlpC, BlpC*Spp*1.1 and BlpC*Spp*2. The latter is identical to BlpC 6A (35) while the former differs from BlpC R6 by one amino acid in the leader peptide sequence (Fig. S5C). Two strains carried other BlpC alleles, BlpC*Spp*1.2, which is identical to BlpC R6 and BlpC*Spp*3 which is unique. Unlike for BlpC, most strains had the same CSP pherotype. Besides CSP6.1 and CSP6.3 which have previously been described in *S. pseudopneumoniae* (36), two new alleles of ComC were found, CSP6.4 and CSP10 (Fig. 6B and S5D).

Genetic determinants of AMR, such as ICEs and Mega-2, as well as phenotypic resistances to penicillin and SXT were mostly associated with clade III, in which 19/20 strains (95,2%) carried at least one genetic element encoding an AMR determinant or have been shown to be resistant to at least one antibiotic (Fig. S6B). A relatively small percentage of strains belonging to clade II (31,6%) were associated with AMR. In general ICE integration sites were shared by closely related strains. 9 of the 11 strains carrying a Mega-2 element are found in a subset of clade III and presence of this element was almost strictly associated with the absence of a plasmid.

### Discussion

Correct identification of SMG strains remains a challenge. While *S. pseudopneumoniae* was originally described as phenotypically different from *S. pneumoniae* using traditional identification methods (1), an increasing number of studies have reported atypical isolates (4, 5, 7, 13). Most likely this is due to the ability of these species to acquire genetic material through natural transformation and to their high genetic relatedness, underlined by our results that nearly 50% of the pan genomes of *S. pneumoniae* and *S. pseudopneumoniae* are shared by both species. Inarguably, the difficulties in identifying *S. pseudopneumoniae* have impaired our understanding of its epidemiology and contribution to human disease. Nonetheless, it was early on found in lower respiratory tract samples and associated with chronic obstructive pulmonary disease (COPD) and exacerbation of COPD (1, 4). While it appears to cause milder infections and to be, at least in some cases, associated with underlying diseases (5, 8), the isolation of *S. pseudopneumoniae* from sterile body sites (7) and from sepsis cases (5) warrants a deeper investigation into this overlooked pathogen. Moreover, as a causative agent is not identified in a significant percentage (≈40%) of LRTI and community-acquired pneumonia cases, both in the community and hospital settings (37), it is a possibility that a fraction of these cases are due to disregarded potential pathogens such as *S. pseudopneumoniae*, that might be discarded as commensals and for which reliable identification methods lack.

In this study we used a collection of suspected *S. pseudopneumoniae* strains isolated from LRTI patients (22). The classification of some of these isolates could only be resolved through WGS and phylogenetic analyses. A thorough comparative genomic analysis allowed us to identify for the first time a genetic marker that is entirely specific to this species, which is a significant advantage compared to other markers which either aim at identifying pneumococci or were found in other SMG species.

Only a surprisingly small percentage (5.6%) of pneumococcal genes known to be differentially regulated in infection- and colonization-relevant conditions were absent from *S. pseudopneumoniae*. Taken together our results indicate that pneumococcal genes important for interaction with its host during invasive disease and cell contact are widespread in *S. pseudopneumoniae*. While all *S. pseudopneumoniae* strains described to date are non-encapsulated, we report here the first isolate encoding and expressing a capsule. The lack of transposase genes on either side of the capsule locus and its higher similarity with the capsular locus of an *S. mitis* strain, argues against its acquisition from a pneumococcal strain. Further studies are needed to understand the biological role of the capsule in *S. pseudopneumoniae*, and to evaluate the prevalence of encapsulated isolates in larger clinical sample collections.

The presence of multiple pneumococcal virulence and colonization factors in the core genome of *S. pseudopneumoniae* confirms earlier observations that many of these genes are found in this species (3, 20). Our results show however that *S. pneumoniae* and *S. pseudopneumoniae* differ in their respective core features. The presence of some of these features, such as pneumolysin, could mark an important difference between *S. pseudopneumoniae* and the more commensal *S. mitis*. Pneumolysin is a core feature of *S. pseudopneumoniae*, whereas it is found in merely 8% of *S. mitis* genomes (data not shown).

Surface-exposed proteins are important players in the successful colonization of its host by the pneumococcus and display a wide variety of function from virulence, to fitness and antibiotic tolerance (38, 39). Although the absence of a capsule in the majority of *S. pseudopneumoniae* strains might be the main reason for its reduced virulence in comparison to pneumococci, the presence of large numbers of surface-exposed proteins could provide an advantage for adhesion and colonization, as was described for NESp (18, 19). In this scenario, the lack of a capsule might avoid restricting the ability of surface-exposed proteins to interact with their ligands on host cells (23). The large number of two-component signalling systems in *S. pseudopneumoniae* might indicate that it is equipped to fine-tune its response to different environmental cues. Taken together, our results reveal that *S. pseudopneumoniae* encodes a large number of novel features that could contribute to virulence, colonization and adaptation.

Our observations reveal a composite scenario of genetic elements in *S. pseudopneumoniae*. The fact that the core genome phylogeny delineates clades that harbour different genetic elements could indicate small differences in their core genome could play a role in the maintenance or exclusion of these elements. Taken together, our observations suggest multiple acquisition events and subsequent clonal expansion of Tn*916*-like ICEs in *S. pseudopneumoniae*. Most of the strains carrying a Mega-2 element are found in a subset of the same clade suggesting its presence is mainly driven through clonal expansion, as was suggested for *S. pneumoniae* (29, 30). Besides genetic determinants of AMR, phenotypic resistances also showed a tight association with a specific lineage. Although no specific virulence factor except for PcpA could be associated with a given clade, it is perhaps worth mentioning that the three septicemia isolates (5) belong to the same phylogenetic clade. In *S. pneumoniae*, longer durations of carriage are associated with increased prevalence of resistance (40). It will be interesting in the future to evaluate the relative virulence of strains belonging to different clades.

Taken together, our results sheds light on the distribution in *S. pseudopneumoniae* of genes known to be important during invasive disease and colonization and reveals the impressive amount of surface-exposed proteins encoded by some strains. While this study does not allow conclusions on the virulence potential of *S. pseudopneumoniae*, our single specific molecular marker for identifying *S. pseudopneumoniae* from other SMG species will be a useful resource for better understanding the clinical importance and epidemiology of this species.

## METHODS

### Bacterial isolates and molecular typing

32 α-hemolytic strains isolated from sputum or nasopharyngeal swabs of lower-respiratory tract infection patients collected during the GRACE study (22) and presenting atypical results in traditional biochemical tests to identify *S. pneumoniae* were included in this study. Isolates were tested for optochin susceptibility as described elsewhere (7) and bile solubility (41) and tested by PCR for pneumococcal markers (*lytA*, *cpsA*, *spn_9802*, 16SrRNA) and by RFLP for pneumococcal-specific signatures (*lytA*, *ply/mly*) (7). BHN880 was serotyped by gel diffusion as described elsewhere (42). MICS to penicillin, sulfamethoxazole-trimethoprim (SXT), erythromycin, clindamycin, tetracycline and levofloxacin were determined using Etests (bioMérieux) and interpreted using the Clinical and Laboratory Standards Insitute (CLSI) guidelines for viridans streptococci (43), except for SXT which was interpreted using the European Committee on Antimicrobial Susceptibility Testing (EUCAST) breakpoints for non-meningitis *S. pneumoniae* isolates (44).

### Whole-genome sequencing, assembly and phylogenetic analysis

Chromosomal DNA was prepared from overnight cultures on blood agar plates using the Genomic DNA Buffer Set and Genomic-tip 100/G (QIAGEN) following manufacturer’s instructions. Long DNA insert sizes were used and Illumina TruSeq HT DNA sample preparation kit was used to prepare libraries. Paired-end reads were generated with read lengths of 250bp. Demultiplexed reads were subjected to adapter removal and were quality trimmed using Trimmomatic (45). The 24 genomes were assembled *de novo* with SPADES (v3.1.1) (46), annotated with PROKKA (v1.11) (47, 48) and deposited in NCBI (XXXX to XXXX). Assembly metrics were calculated with QUAST 4.5.4 (48). kSNP 3.1 (49) was used to generate a SNP-based phylogenetic tree, using NCBI genomes of *S. pseudopneumoniae* (n=38), *S. mitis* (n=36), completed genomes of *S. pneumoniae* (n=39), *S. oralis* (n=1), *S. infantis* (n=1), non-typable *S. pneumoniae* recently identified as *S. pseudopneumoniae* (n=8) (12) and our 24 LRTI isolates. The optimum K-mer value of 19 estimated from Kchooser and a consensus parsimony tree based on all the SNPs generated by kSNP was used (49). The phylogenetic tree was visualized in MEGA7 (50).

### Pan genome analysis, construction of SPPN species tree and identification of virulence factors

The pan-genome analysis of orthologous gene clusters, species trees and their respective gene trees were analyzed using panX (51) for the 39 completed strains of *S. pneumoniae* [pan:SPN], 44 *S. pseudopneumoniae* [pan:SPPN] and both the species [pan: SPPN-SPN] with the default cut-off values. pan:SPPN analysis resulted in 885 core genes (strict core; 100% present in all the strains) and the core-genome tree/species tree for the SPPN species was constructed based on the core-genome SNPs including only single copy core genes (n=793). Using pan:SPPN-SPN, all COGs were queried for the *S. pneumoniae* locus tags corresponding to the 356 virulence genes (23) and 92 well studied pneumococcal genes (26) listed in Tables S4 and S5. Additionally, the proteins listed in Table S5 were analyzed using a 70% length cutoff to score proteins as present; conservation of synteny with was confirmed for all proteins. Genetic loci of proteins scored as absent were manually checked for contig breaks and pseudogenes.

### Molecular markers and PCR assay

30 unique gene clusters present in the 44 *S. pseudopneumoniae* genomes and absent from the 39 *S. pneumoniae* genomes were filtered from the pan genome analysis and blasted against all NCBI genomes. The 44 nucleotide sequence of the two unique ORFs (SPPN_RS10375 and SPPN_RS06420) were aligned using the ClustalW algorithm in Geneious version 10.1.3 (https://www.geneious.com) with default parameters (Gap open cost = 15, Gap extend cost = 6.66). The upstream (70 bp) and downstream (329 bp) intergenic regions of SPPN_RS10375 were included. Primers SPPN_RS10375F (5’-CTAATTGCTACTGCTATTTCCGGTG-3’) and SPPN_RS10375R (5’-CTGATACCTGCAACAAAAATCGAAG-3’) were designed in conserved regions. PCR was performed using PHUSION Flash High-Fidelity PCR Master Mix (ThermoFisher) following manufacturer’s instructions and with an annealing temperature of 50°C. 1 ul of lysate prepared by resuspending 2-3 isolated colonies in 100 ul TE containing 0.1% Triton and incubating at 98°C for 5 min in a dry bath was used as template in each PCR reaction. PCR products were run on a 1.2% agarose gel stained with GelRed (Biotium).

### Analysis of the capsular loci

Homologues of *cpsA/wzg* were searched for in pan:SPPN_SPN using gene family SP_RS01690. The locus was then subsequently checked manually for the presence of the complete locus [BHN880_01411 - BHN880_01431]. The retrieved *cps* locus was blasted (Blastn) to identify the closest homologs. Pairwise alignment with the serotype 5 reference locus (CR931637.1) (24) and *S. mitis* 21/39 (AYRR01000010.1) *cps* locus was performed using Easyfig (52).

### *In silico* identification of new putative virulence features

The proteins from all the pseudopneumoniae genomes (n=44) were concatenated to build the SPPN protein database. Using the NCBI Batch CD-Search tool (53), the SPPN protein database was queried for the presence of the conserved choline-binding domain COG5263 and the peptidase_M26 domain pfam07580/ cl06563 to identify the novel choline-binding proteins (CBPs) and zinc-metalloproteases (ZMPs) respectively. Two-component signal transduction systems (TCSs) were identified in a similar way by searching for the HATPase domain of the histidine kinase protein (cd00075/smart00387/pfam02518) with immediately preceded or followed by a DNA-binding regulator possessing the signal-receiver domain, cd00156.

### *In silico* identification of AMR determinants, plasmids and phages

The 44 genomes were screened in Resfinder 3.0 (54) for acquired antibiotic resistance (AMR) genes (90% identity threshold, minimum length of 60%). Chromosomal genes flanking Tn*916*-like ICEs were defined by using BLASTn to retrieve the loci in strain IS7493 (NC_015875.1) of the genes located immediately upstream the integrase and immediately downstream *orf24* of Tn*5251* (FJ711160.1). Genome assemblies were queried for genes associated with known *S. pneumoniae* and *S. mitis* phages, and the *S. pseudopneumoniae* plasmid pDRPIS7493 (NC_015876.1) (Table S10). Phage sequences were manually analyzed and deemed full-length if they started with an integrase gene, ended with a lytic amidase and were ≥30 kb in length.

## ACKNOWLEDGEMENTS

LRTI samples were collected as part of the GRACE (Genomics to combat resistance against antibiotics in CA-LRTI in Europe) project. We thank the GPs, the GRACE study team, and the patients for taking part in this study. We thank the European commission for the financial support of the GRACE project. We also thank Ingrid Andersson, Christina Johansson, Gunnel Möllerberg, Eva Morfeldt, and Jessica Darenberg at the Swedish Institute for Infectious Disease Control for excellent technical assistance. We acknowledge support from the National Genomics Infrastructure in Stockholm funded by Science for Life Laboratory, the Knut and Alice Wallenberg Foundation and the Swedish Research Council, and SNIC/Uppsala Multidisciplinary Center for Advanced Computational Science for assistance with massively parallel sequencing and access to the UPPMAX computational infrastructure. This work was partially supported by ONEIDA project (LISBOA-01-0145-FEDER-016417) co-funded by FEEI - “Fundos Europeus Estruturais e de Investimento” from “Programa Operacional Regional Lisboa 2020” and by national funds from Fundação para a Ciência e a Tecnologia, Portugal.

